# A species-wide genetic atlas of antimicrobial resistance in *Clostridioides difficile*

**DOI:** 10.1101/2021.06.14.448453

**Authors:** Korakrit Imwattana, César Rodríguez, Thomas V. Riley, Daniel R. Knight

## Abstract

Antimicrobial resistance (AMR) plays an important role in the pathogenesis and spread of *Clostridioides difficile* infection (CDI), the leading healthcare-related gastrointestinal infection in the world. An association between AMR and CDI outbreaks is well documented, however, data is limited to a few ‘epidemic’ strains in specific geographical regions. Here, through detailed analysis of 10,330 publicly-available *C. difficile* genomes from strains isolated worldwide (spanning 270 multilocus sequence types (STs) across all known evolutionary clades), this study provides the first species-wide snapshot of AMR genomic epidemiology in *C. difficile*. Of the 10,330 *C. difficile* genomes, 4,532 (43.9%) in 89 STs across clades 1 – 5 carried at least one genotypic AMR determinant, with 901 genomes (8.7%) carrying AMR determinants for three or more antimicrobial classes (multidrug-resistant, MDR). No AMR genotype was identified in any strains belonging to the cryptic clades. *C. difficile* from Australia/New Zealand had the lowest AMR prevalence compared to strains from Asia, Europe and North America (p<0.0001). Based on the phylogenetic clade, AMR prevalence was higher in clades 2 (84.3%), 4 (81.5%) and 5 (64.8%) compared to other clades (collectively 26.9%) (p<0.0001). MDR prevalence was highest in clade 4 (61.6%) which was over three times higher than in clade 2, the clade with the second-highest MDR prevalence (18.3%). There was a strong association between specific AMR determinants and three major epidemic *C. difficile* STs: ST1 (clade 2) with fluoroquinolone resistance (mainly T82I substitution in GyrA) (p<0.0001), ST11 (clade 5) with tetracycline resistance (various *tet*-family genes) (p<0.0001) and ST37 (clade 4) with macrolide-lincosamide-streptogramin B (MLS_B_) resistance (mainly *ermB*) (p<0.0001) and MDR (p<0.0001). A novel and previously overlooked *tetM*-positive transposon designated Tn*6944* was identified, predominantly among clade 2 strains. This study provides a comprehensive review of AMR in the global *C. difficile* population which may aid in the early detection of drug-resistant *C. difficile* strains, and prevention of their dissemination world-wide.

**Impact statement:** Utilising a publicly-available database of 10,330 sequence reads, this study provides the first species-wide evaluation of genotypic AMR in *C. difficile*. It reports the most common AMR determinants and their genomic neighbourhood, associations between important genotypes and specific strains or geographical regions, and rare AMR genotypes that may have been missed in earlier studies.

**Data summary:** This study utilises publicly available raw sequence reads available at the NCBI Sequence Read Archive (SRA) as of January 2020. The details of all genomes are available in the **Supplementary Data** (10.6084/m9.figshare.14623533).

## Introduction

Antimicrobial resistance (AMR) is one of the biggest threats to modern medicine. Without focused interventions and collaborations across all government sectors, AMR could be responsible for an estimated 10 million deaths and the loss of up to US$210 trillion of annual global income by 2050 (1). The US Centers for Disease Control and Prevention (CDC) reported on AMR health threats in 2013 (2), with an update in 2019 (3), highlighting organisms with the highest AMR burden and threat (3).

*Clostridioides* (*Clostridium*) *difficile* infection (CDI) causes major gastrointestinal illness worldwide (4), responsible for as many as 14,000 deaths annually in the US (2). *C. difficile* has been classified by the CDC as an urgent threat, the highest threat level, in both the 2013 and 2019 CDC reports, responsible for the highest number of annual deaths among the pathogens listed (2, 3). In contrast to other pathogens, AMR in *C. difficile* has some unique features. AMR leads to difficulties in treating infections (5), and although the treatment of CDI is also a challenge (6), the challenge is not due to AMR *per se* as resistance to antimicrobials used for the treatment of CDI remains rare (7). Instead, AMR plays a significant role in the pathogenesis and spread of CDI (8).

Using multi-locus sequence typing (MLST), the population of *C. difficile* can be divided into five major clades (C1 – C5) and three smaller cryptic clades. The three cryptic clades are extremely divergent (**Figures 1 and 2A**) and likely represent independent species or subspecies (9). To date, extensive studies have been conducted on the role of AMR in the emergence and spread of two epidemic PCR ribotypes (RTs) of *C. difficile* RTs 027 and 078, which correspond to multilocus sequence types (STs) 1 and 11, respectively (10–12). A few studies have focused also on *C. difficile* RT 017 (ST 37) (13), a third epidemic lineage (14), which shows a high prevalence of resistance to many antimicrobial classes (8). Although these studies provided insights on how AMR impacts the spread of *C. difficile*, they are limited to a few strain types in specific geographical regions, and there has not been any study of AMR prevalence in the species-wide population of *C. difficile*. Here, through detailed analysis of 10,330 publicly-available genomes from *C. difficile* isolated worldwide, we provide the first species-wide snapshot of AMR genomic epidemiology in *C. difficile*.

**Figure 1 –.**
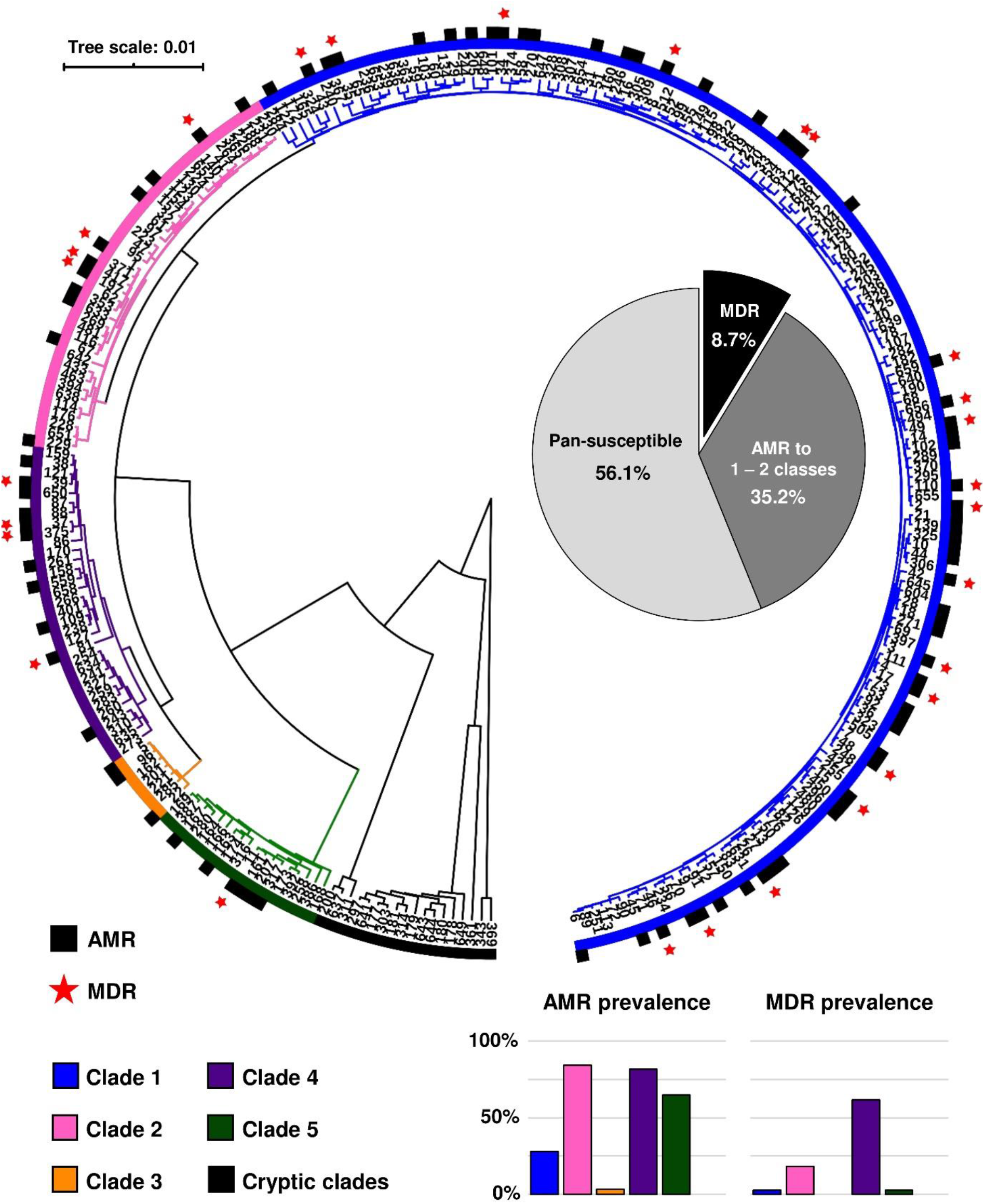
Distribution of resistant and multidrug-resistant *C. difficile*. The UPGMA phylogenetic tree represents a total of 270 STs included in this study. The black sections indicate that at least one strain in the ST had acquired resistance (AMR) to at least one antimicrobial class. The red stars indicate that at least one strain in the ST was MDR (i.e. had acquired resistance to at least three antimicrobial classes). The pie chart in the middle shows the overall prevalence of MDR *C. difficile* (black), *C. difficile* resistance to 1-2 antimicrobial classes (dark grey) and pan-susceptible *C. difficile* (light grey) among 10,330 *C. difficile* strains. The bar charts below show the prevalence of resistant and MDR strains in each clade.

**Figure 2 –.**
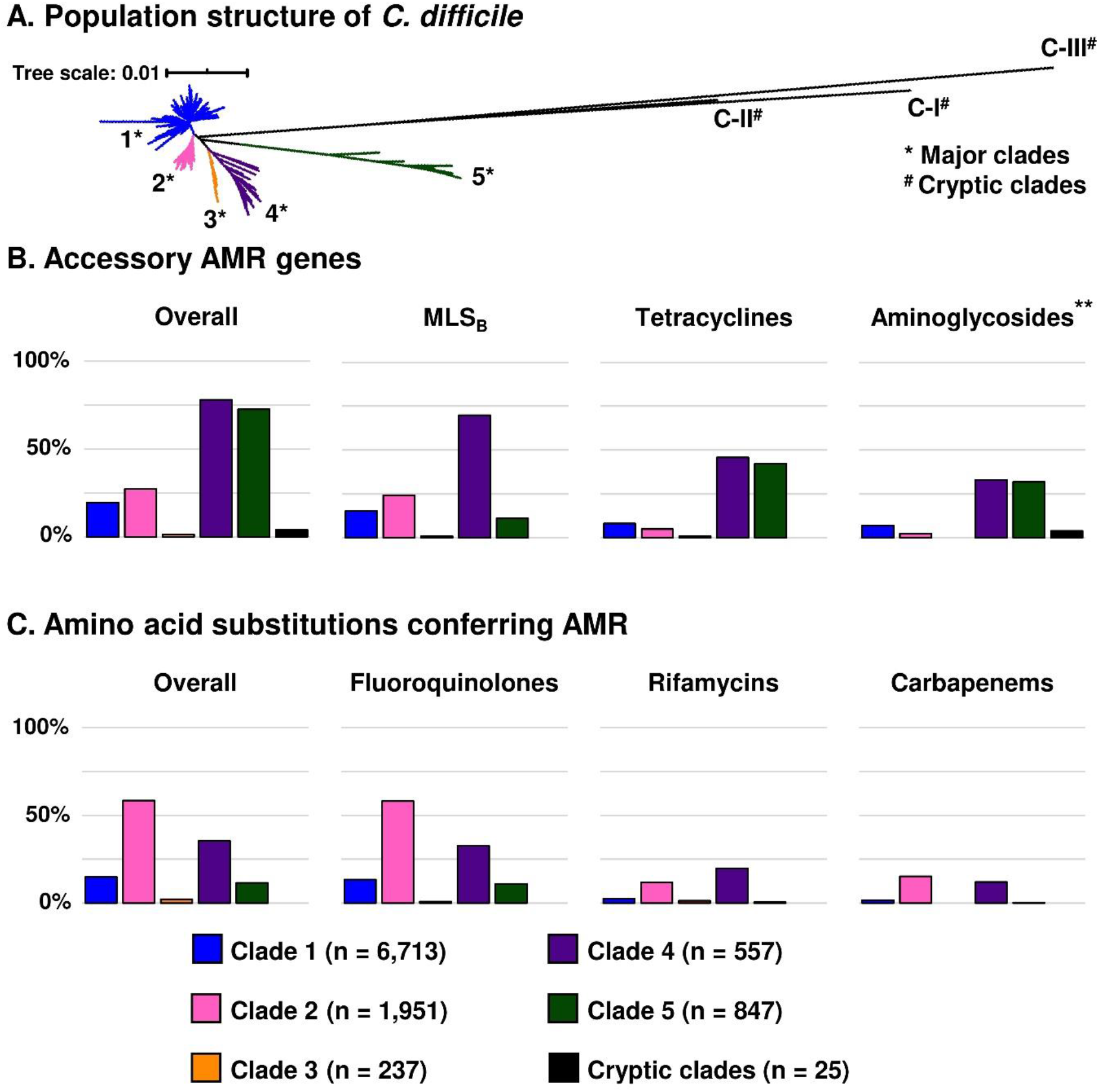
Summary of antimicrobial resistance genotype of *C. difficile*. **(A)** For evolutionary context, a neighbour-joining phylogeny based on MLST shows the global population structure of *C. difficile*. **(B)** The prevalence of *C. difficile* strains harbouring accessory AMR genes across different clades (leftmost) and the prevalence of resistance to important antimicrobial classes conferred mainly by accessory AMR genes. The presence of an aminoglycoside resistance gene (**) does not contribute to the definition of MDR *C. difficile*. **(C)** The prevalence of *C. difficile* strains having significant amino acid substitutions associated with AMR across different clades (leftmost) and the prevalence of resistance to important antimicrobial classes conferred mainly by amino acid substitution.

## Materials and Methods

### Genome collection and de-replication of clonal strains

The starting point for this analysis was an international collection of 12,098 *C. difficile* Illumina paired-end sequence reads sourced from the NCBI Sequence Read Archive (SRA, https://www.ncbi.nlm.nih.gov/sra/) in January 2020. All sequence reads were screened for contamination using Kraken2 v2.0.8-beta and only reads with >85% of sequences classified as *C. difficile* were included. MLST was confirmed on these raw sequence reads by SRST2 v0.2.0 with the database available on PubMLST (https://pubmlst.org/organisms/clostridioides-difficile) as previously described (9, 15). This dataset comprised a total of 270 STs spanning the eight currently described evolutionary clades with a relatively high number of reads from epidemic strains, particularly STs 1 (C2; n=2,532), 11 (C5; n=1,185) and 37 (C4; n=786), many of which were likely to be clonal. To adjust for this strain selection bias, pairwise average nucleotide identity (ANI) of reads from these three STs, as well as ST 2 (n=1,153), the most common strain in C1, were compared using the Sketch algorithm included in BBtools (https://sourceforge.net/projects/bbmap/). Reads with an ANI of 99.98% or higher were considered to be clonal and only one genome from each clonal complex was included in the final analysis. Based on a small dataset of 240 *C. difficile* reads (28,680 possible pairs, 531 of which were clonal pairs), this cut-off point had a sensitivity of 70.1% and a specificity of 76.8% for the detection of clonal strains as defined by Didelot *et al* (data not shown) (16). The 10,330 reads remaining in the dataset are summarised in **Table 1**.

**Table 1.** *C. difficile* strains in the de-replicated NCBI database (January 2020).

| *C. difficile* clade | Number of genomes (%) | Most prevalent STs |
| --- | --- | --- |
| C1 | 6,713 (65.0%) | ST 2 (9.2%)*<br>ST 8 (6.0%)*<br>ST 3 (5.4%)*<br>ST 42 (4.1%)*<br>ST 6 (3.2%)*<br>ST 44 (2.5%)*<br>ST 14 (2.4%)* |
| C2 | 1,951 (18.9%) | ST 1 (16.6%)*<br>ST 41 (0.8%) |
| C3 | 237 (2.3%) | ST 5 (2.0%)<br>ST 22 (0.2%) |
| C4 | 557 (5.4%) | ST 37 (4.3%)*<br>ST 39 (0.2%) |
| C5 | 847 (8.2%) | ST 11 (7.6%)*<br>ST 167 (0.1%) |
| Cryptic clades | 25 (0.2%) | ST 361 (<0.1%)<br>ST 177 (<0.1%) |
| **Total** | **10,330** | - |

Note: \* Ten most prevalent sequence types (STs) in this dataset.

### *Identification of multidrug-resistant* C. difficile

Multidrug-resistant (MDR) *C. difficile* in this study refers to *C. difficile* strains with genotypic AMR determinants (both accessory genes and mutations in chromosomal genes) for at least three of the following antimicrobial classes: carbapenems, fluoroquinolones, glycopeptides (vancomycin), nitroimidazoles (metronidazole), oxazolidinones (linezolid), macrolide-lincosamide-streptogramin B (MLS_B_), phenicols, rifamycins, tetracyclines and sulfa-containing agents. Resistance determinants for aminoglycosides and cephalosporins were excluded from this definition as *C. difficile* is intrinsically resistant to these agents (17, 18).

### Detection of accessory AMR genes and associated transposons

To detect the presence of accessory AMR genes, raw sequence reads were interrogated against ResFinder/ARGannot databases, with an addition of two newly-characterised AMR genes found in *C. difficile, erm*(52) and *mefH*, using SRST2 with default settings (15, 19–21). These databases contain over 500 different genes conferring resistance to 15 different antimicrobial classes, covering all AMR genes known to be carried by the *C. difficile* population analyzed so far (19, 20). The spectrum of *β*-lactamase enzymes detected was confirmed against the CARD 2020 database (22). To further characterise the genomic context of the most common accessory AMR genes, *C. difficile* strains with *ermB, tetM* and *tet44* genes were interrogated using SRST2 against a database of *C. difficile* transposons carrying *ermB* (Tn*5398* [GenBank accession AF109075.2], Tn*6189* [MK895712.1], Tn*6194* [HG475346.1], Tn*6215* [KC166248.1] and Tn*6218* [HG002387.1]), *tetM* (Tn*916* [U09422.1], Tn*5397* [AF333235.1] and Tn*6190* [FN665653]) and *tet44* (Tn*6164* [FN665653]) (23, 24) with 80% minimum coverage and 10% maximum divergence (15), corresponding with 72% minimum nucleotide identity (NI).

To detect the presence of a plasmid conferring metronidazole resistance (pCD-METRO) (25), a custom database was created consisting of all eight coding sequences (CDS) of pCD-METRO. SRST2 was used with default settings on all sequence reads against this customised database (15). The 23 *C. difficile* genomes from the original study (25) were included in the analysis and used to evaluate the accuracy of the database.

### Detection of amino acid substitutions conferring AMR

All genomes were screened for known point mutations in *gyrA, gyrB, rpoB, pbp1* and *pbp3* genes using customised databases in SRST2. The reference sequences for these genes were obtained from the PubMLST database (https://pubmlst.org/organisms/clostridioides-difficile/) as well as reference *C. difficile* genomes (CD630 [C1, GenBank accession AM180355], CD196 [C2, FN538970], M68 [C4, FN668375] and M120 [C5, FN665653]). *C. difficile* strains were categorized as resistant to an antimicrobial if they carried a gene allele with at least one significant point mutation listed in **Table 2** (23, 26, 27).

**Table 2.** Summary of known non-synonymous chromosomal point mutations conferring AMR.

| Protein | Substitution | Clade distribution* |  |  |  |  |  | Comment |
| --- | --- | --- | --- | --- | --- | --- | --- | --- |
|  |  | C1 | C2 | C3 | C4 | C5 | Cryptic |  |
| Fluoroquinolone resistance |  |  |  |  |  |  |  |  |
| GyrA | Val43Asp | ○ | ○ | ○ | ○ | ○ | ○ | Absent in this dataset |
|  | Asp71Val | ● | ○ | ○ | ● | ○ | ○ | Found in < 10 strains in this dataset |
|  | Asp81Asn | ● | ○ | ○ | ○ | ○ | ○ | Found in < 10 strains in this dataset |
|  | Thr82Ile | ● | ● | ● | ● | ● | ○ | Most common substitution |
|  | Thr82Val | ● | ○ | ○ | ○ | ● | ○ | Found in < 10 strains in this dataset |
|  | Ala118Thr | ● | ○ | ○ | ○ | ● | ○ | Found in < 10 strains in this dataset |
|  | Ala384Asp | ● | ○ | ○ | ○ | ○ | ○ | Found in < 10 strains in this dataset |
| GyrB | Arg377Gly | ○ | ○ | ○ | ○ | ○ | ○ | Absent in this dataset |
|  | Asp426Asn | ● | ● | ○ | ● | ○ | ○ | Most common substitution |
|  | Asp426Val | ● | ○ | ○ | ● | ○ | ○ | Mostly found in clade 4 *C. difficile* |
|  | Arg447Lys | ● | ● | ○ | ○ | ● | ○ |  |
|  | Glu466Val | ○ | ○ | ○ | ○ | ● | ○ | Found in < 10 strains in this dataset |
| Rifamycin resistance |  |  |  |  |  |  |  |  |
| RpoB | Asp492Asn | ○ | ○ | ○ | ○ | ○ | ○ | Absent in this dataset |
|  | Asp492Val | ○ | ○ | ○ | ○ | ○ | ○ | Absent in this dataset |
|  | His502Asn | ● | ● | ○ | ● | ○ | ○ |  |
|  | His502Arg | ○ | ○ | ○ | ○ | ○ | ○ | Absent in this dataset |
|  | His502Leu | ○ | ○ | ○ | ○ | ○ | ○ | Absent in this dataset |
|  | His502Tyr | ● | ○ | ○ | ○ | ○ | ○ | Found in < 10 strains in this dataset |
|  | Arg505Lys | ● | ● | ● | ● | ● | ○ | Most common substitution |
|  | Ser550Phe | ● | ● | ○ | ○ | ● | ○ | Found in < 10 strains in this dataset |
|  | Ser550Tyr | ● | ● | ○ | ● | ○ | ○ | Found in < 10 strains in this dataset |
| Fidaxomicin resistance |  |  |  |  |  |  |  |  |
| RpoB | Gln1073Arg | ○ | ○ | ○ | ○ | ○ | ○ | Absent in this dataset |
| Carbapenem resistance |  |  |  |  |  |  |  |  |
| Pbp1 | Leu543His | ● | ○ | ○ | ○ | ○ | ○ |  |
|  | Ala555Thr | ● | ● | ○ | ● | ● | ○ | Most common substitution |
| Pbp3 | Tyr721Ser | ● | ○ | ○ | ● | ○ | ○ |  |

Note: \* Based on significant findings in this study. Solid circles refer to the presence of the substitution in the clade.

### Assessment of AMR prevalence in different geographical areas

Data on geographical regions of isolation was available for 6,227 (60.3%) *C. difficile* strains: Asia (n = 355), Europe (n = 3,548), North America (n = 2,212) and Australia/New Zealand (n = 112). The clade distribution was notably different in these regions (**Table 3**). Thus, multiple logistic regression analyses were performed using R to assess the clade-adjusted AMR prevalence for major antimicrobial classes (MLS_B_, tetracyclines, fluoroquinolones and rifamycins), as well as MDR prevalence. From the initial analysis, the overall AMR prevalence was lowest in strains from Australia/New Zealand. Thus, they were used as the reference group in this analysis.

**Table 3.** Clade distribution in 4 major geographical regions.

| Region | Clade |  |  |  |  |  |
| --- | --- | --- | --- | --- | --- | --- |
|  | C1 | C2 | C3 | C4 | C5 | Cryptic |
| Asia | 76.6% | 3.4% | 3.9% | 15.2% | 0.6% | 0.3% |
| Europe | 74.4% | 11.0% | 3.4% | 2.7% | 8.3% | 0.2% |
| North America | 68.9% | 24.7% | 0.1% | 2.1% | 4.0% | 0.2% |
| Australia/New Zealand | 39.3% | 26.8% | 1.8% | 3.6% | 28.6% | 0.0% |

## Results

### Summary of AMR and MDR prevalence

Of the 10,330 *C. difficile* genomes evaluated, 4,532 (43.9%) contained acquired resistance genes for at least one antimicrobial class, with 89 STs across 5 major clades having at least one resistant strain (**Figure 1**). A total of 901 strains (8.7%) across 28 STs harboured resistance determinants for three or more antimicrobial classes and were therefore classified as MDR. Based on resistance prevalence, *C. difficile* could be divided into clades with an overall resistance prevalence of ≥ 50%, which included C2, C4 and C5, each of which contained an epidemic ST (ST 1 in C2, ST 37 in C4 and ST 11 in C5), and clades with an overall resistance prevalence of < 50%, which included C1 and C3, as well as all three cryptic clades. The prevalence of MDR *C. difficile* was highest in C4 *C. difficile* (61.6% compared to an overall 5.7% in other clades), over three times higher than in C2 which had the second-highest prevalence of MDR strains (18.3%). The overall resistance prevalence of important antimicrobial classes is shown in **Figure 2**.

### AMR prevalence in different geographical regions

**Figure 3** shows the results of logistic regression analyses of the clade-adjusted AMR and MDR prevalence compared to strains from Australia/New Zealand as the reference. Overall, strains from Asia, Europe and North America all had higher AMR prevalence (p< 0.0001). The difference in AMR prevalence was most pronounced for fluoroquinolones, where the prevalence of substitution associated with fluoroquinolone resistance (FQR) in the three continents (collectively 1,491/6,115; 24.4%) was estimated to be at least nine times higher than in Australia/New Zealand (3/112; 2.7%). In Asia, Europe and North America, AMR prevalence was not significantly different, with AMR prevalence in Asia (99/355; 27.9%) marginally higher than in Europe (814/3,548; 22.9%) and North America (578/2,212; 26.1%).

### Fluoroquinolone resistance

Overall, 2,959 *C. difficile* strains (28.6%) carried known DNA gyrase substitutions associated with fluoroquinolone resistance (FQR). The prevalence of FQR was highest in clade C2 (82.3%), followed by C4 (53.1%). Most resistance was conferred by point substitutions solely within the GyrA subunit of the enzyme (2,771 strains, 93.7% of all resistant strains), followed by point substitutions solely within the GyrB subunit (104 strains, 3.5%). Only 84 strains (2.8%) had substitutions on both gyrase subunits. The prevalence of GyrB subunit substitution (both alone and in addition to GyrA substitution) was highest in C4 (10.6%). The most common GyrA substitution was Thr82Ile (found in 2,843 strains, 99.6% of strains with GyrA substitution) and the most common GyrB substitution was Asp426Asn (131 strains, 69.7% of strains with GyrB substitution), followed by Asp426Val (44 strains, 23.4%), the latter was almost exclusive to C4 (40/44 strains, 90.9%). Interestingly, a Ser416Ala substitution, a polymorphism that does not confer resistance, was found in a majority of C5 (825 strains, 94.9%) and cryptic clades (20 strains, 80.0%), but in only one clade C1 strain and none of the other major clades.

**Figure 3.**
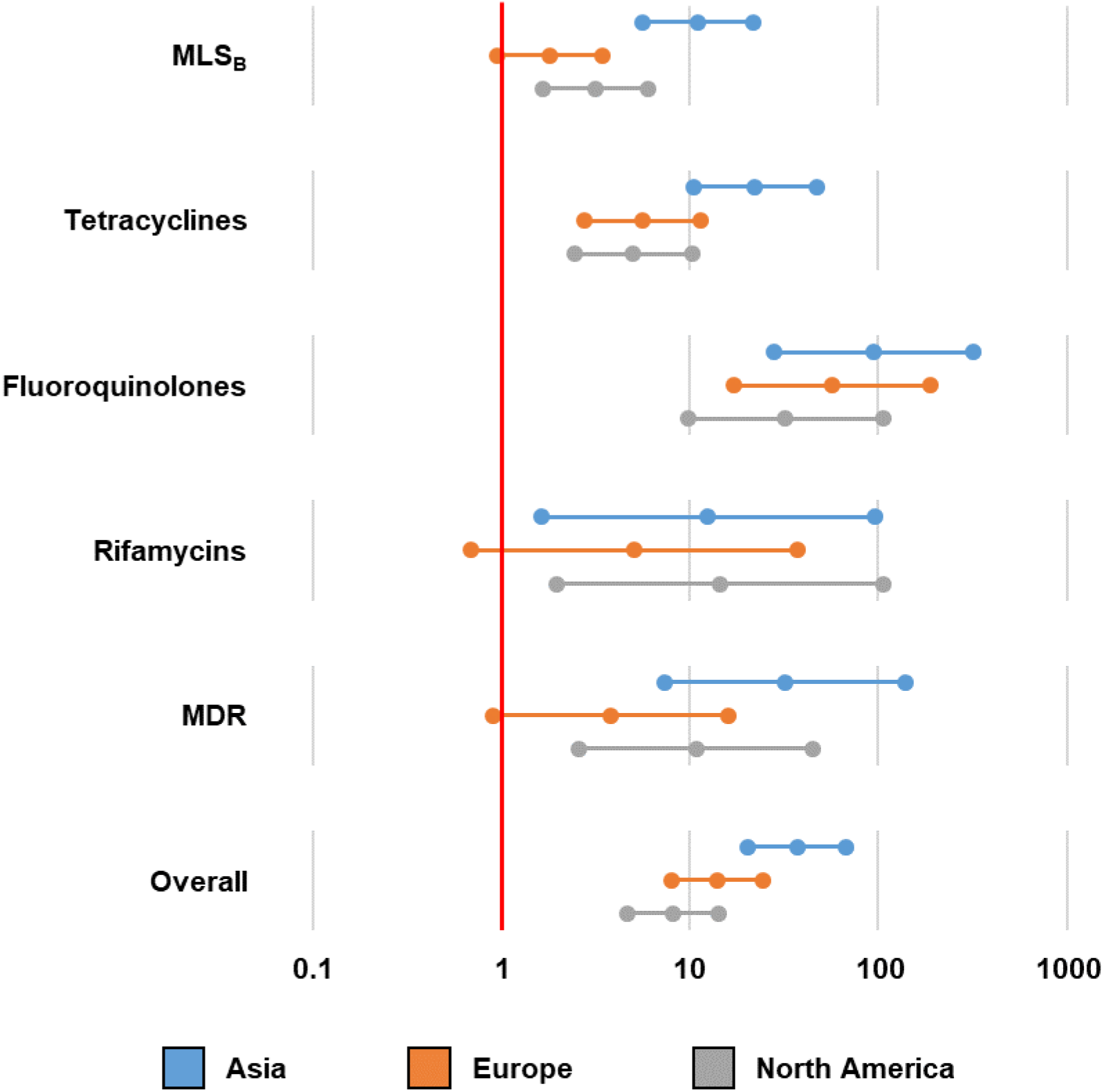
Difference in AMR prevalence in different geographical regions. Multiple logistic regression analyses were performed to compare the clade-adjusted AMR prevalence in four regions (Asia, Europe, North America and Australia/New Zealand). The Forest plot represents the estimated AMR prevalence in each continent compared to Australia/New Zealand.

### MLS_B_ resistance

**Table 4** summarises the major genotypic determinants for MLS_B_ antimicrobials detected in our survey. The most common determinants were *ermB* (1,775 strains, 17.2%) followed by *erm*(52) (145 strains, 1.4%) and *ermG* (25 strains, 0.2%). The *erm* class genes, which methylate 23S rRNA and prevent the binding of MLS_B_ antimicrobials, are associated with high-level resistance to all MLS_B_ antimicrobials, as shown by high-level resistance to both clindamycin and erythromycin (28). The most common non-*erm* genes were *mefH* (156 strains, 1.5%), *mefA* (24 strains, 0.2%), *msrD* (21 strains, 0.2%) and *lnuC* (17 strains, 0.2%). In total, 1,979 *C. difficile* strains (19.2%) across 65 STs (23.9%) in five major clades carried acquired MLS_B_ resistance determinants.

**Table 4.** Summary of resistance determinants for MLS_B_ antimicrobials.

| Gene | Clade distribution [N (%)] |  |  |  |  |  | Overall |
| --- | --- | --- | --- | --- | --- | --- | --- |
|  | C1 | C2 | C3 | C4 | C5 | Cryptic |  |
| *ermB* | 953<br>(14.2%) | 421<br>(21.6%) | 0<br>(0.0%) | 328<br>(58.9%) | 73<br>(8.6%) | 0<br>(0.0%) | 1,776<br>(17.2%) |
| Tn5398 | 168<br>(2.5%) | 1<br>(0.1%) | 0<br>(0.0%) | 0<br>(0.0%) | 1<br>(0.1%) | 0<br>(0.0%) | 170<br>(1.6%) |
| Tn6189 | 259<br>(3.9%) | 104<br>(5.3%) | 0<br>(0.0%) | 44<br>(7.9%) | 17<br>(2.0%) | 0<br>(0.0%) | 424<br>(4.1%) |
| Tn6194 | 204<br>(3.0%) | 270<br>(13.8%) | 0<br>(0.0%) | 268<br>(48.1%) | 46<br>(5.4%) | 0<br>(0.0%) | 788<br>(7.6%) |
| Tn6215 | 106<br>(1.6%) | 0<br>(0.0%) | 0<br>(0.0%) | 0<br>(0.0%) | 2<br>(0.2%) | 0<br>(0.0%) | 108<br>(1.0%) |
| Tn6218 | 200<br>(3.0%) | 4<br>(0.2%) | 0<br>(0.0%) | 10<br>(1.8%) | 2<br>(0.2%) | 0<br>(0.0%) | 216<br>(2.1%) |
| Unknown | 16<br>(0.2%) | 42<br>(2.2%) | 0<br>(0.0%) | 6<br>(1.1%) | 5<br>(0.6%) | 0<br>(0.0%) | 69<br>(0.7%) |
| Other *erm* genes | 86<br>(1.3%) | 17<br>(0.9%) | 1<br>(0.4%) | 66<br>(11.8%) | 4<br>(0.5%) | 0<br>(0.0%) | 175<br>(1.7%) |
| Non-*erm* genes | 76<br>(1.1%) | 104<br>(5.3%) | 1<br>(0.4%) | 22<br>(3.9%) | 18<br>(2.1%) | 0<br>(0.0%) | 222<br>(2.1%) |

Among *ermB*-positive strains, known *ermB*-carrying transposons were identified in 1,706 strains (96.5%) (range, 77.6% – 100.0% NI). Transposon diversity was highest in C1 (**Table 4**). The most common *ermB*-positive transposon was Tn*6194* (788 strains, 44.4%; 81.9% – 100.0% NI), followed by Tn*6189* (424 strains, 23.9%; 77.6% – 99.9% NI) and Tn*6218* (216 strains, 12.2%; 85.3% – 100.0% NI). Tn*5398*, which contains two copies of the *ermB* gene, was found in 170 strains (9.6%; 81.2% – 100.0% NI), most of which belonged to clade C1 (168 strains, 98.8%).

### Tetracycline resistance

**Table 5** summarises the genotypic determinants found for tetracyclines. The most common tetracycline resistance determinant was *tetM* (1,447 strains, 14.0%), followed by *tet40* (214 strains, 2.1%) and *tet44* (125 strains, 1.2%). These three genes encode ribosomal protection proteins which prevent the binding of tetracyclines to 16S rRNA. In total, 1,645 *C. difficile* strains (15.9%) across 68 STs (25.0%) in five major clades carried at least one *tet* gene, with 333 strains (3.2%) carrying more than one gene, 271 of which (81.4%) belonged to clade C5. Five ST11 *C. difficile* strains (C5) carried four different *tet* genes, the highest number of *tet* genes per genome in this dataset. Interestingly, *tet40* and *tet44* were almost exclusively found in clade C5 *C. difficile* (94.9% and 98.4% of *tet40-* and *tet44-*positive *C. difficile* belonged to C5, respectively).

**Table 5.** Summary of resistance determinants for tetracyclines.

| Gene | Clade distribution [N (%)] |  |  |  |  |  | Overall |
| --- | --- | --- | --- | --- | --- | --- | --- |
|  | C1 | C2 | C3 | C4 | C5 | Cryptic |  |
| *tetM* | 457<br>(6.8%) | 128<br>(6.6%) | 0<br>(0.0%) | 402<br>(72.2%) | 460<br>(54.3%) | 0<br>(0.0%) | 1,447<br>(14.0%) |
| Tn916 | 146<br>(2.2%) | 25<br>(1.3%) | 0<br>(0.0%) | 95<br>(17.1%) | 298<br>(35.2%) | 0<br>(0.0%) | 564<br>(5.5%) |
| Tn5397 | 215<br>(3.2%) | 1<br>(0.1%) | 0<br>(0.0%) | 1<br>(0.2%) | 8<br>(0.9%) | 0<br>(0.0%) | 225<br>(2.2%) |
| Tn6190 | 7<br>(0.1%) | 2<br>(0.1%) | 0<br>(0.0%) | 297<br>(53.3%) | 150<br>(17.7%) | 0<br>(0.0%) | 456<br>(4.4%) |
| Tn6944 | 52<br>(0.8%) | 97<br>(5.0%) | 0<br>(0.0%) | 6<br>(1.1%) | 1<br>(0.1%) | 0<br>(0.0%) | 156<br>(1.5%) |
| Unknown | 37<br>(0.6%) | 3<br>(0.2%) | 0<br>(0.0%) | 3<br>(0.5%) | 3<br>(0.4%) | 0<br>(0.0%) | 46<br>(0.4%) |
| *tet44* | 2<br>($<0.1\%$) | 0<br>(0.0%) | 0<br>(0.0%) | 0<br>(0.0%) | 123<br>(14.5%) | 0<br>(0.0%) | 125<br>(1.2%) |
| Tn6164 | 2<br>($<0.1\%$) | 0<br>(0.0%) | 0<br>(0.0%) | 0<br>(0.0%) | 123<br>(14.5%) | 0<br>(0.0%) | 125<br>(1.2%) |
| Other *tet* genes | 129<br>(1.9%) | 12<br>(0.6%) | 2<br>(0.8%) | 14<br>(2.5%) | 336<br>(39.7%) | 0<br>(0.0%) | 493<br>(4.8%) |

Known *tetM*-positive transposons and their variants were detected in 1,245 (86.0%) *tetM*-positive *C. difficile* (78.0 – 100.0% NI). Transposon diversity was highest in clade C1 (**Table 5**). The most common transposons were Tn*916* (564 strains, 39.0%; 83.3% – 100.0% NI) and Tn*6190* (456 strains, 31.5%; 81.5% – 100.0% NI). In contrast to the prevalence of *ermB*-positive transposons above, the distribution of *tetM*-positive transposons was different in clades C2, C4 and C5 (**Figure 4A**). Known *tetM*-positive transposons could not be identified in 78.1% of *tetM*-positive clade C2 *C. difficile* (100/128 strains). Analysis of the assembled genome of ST1 strain C00008355, a clinical isolate from the UK [SRA accession ERR347593], showed that the *tetM* gene was located on a 9,013 bp element with an overall 37.1% GC which did not match any transposons in the NCBI database or published literature (**Figure 4B**). The annotated sequence of this novel Tn, designated Tn*6944* by the Liverpool transposon repository (29), was submitted to GenBank and is available in the DDBJ/ENA/GenBank databases under the accession number BK013348. Besides *tetM*, Tn*6944* also carries *mefH* which encodes a macrolide efflux protein (21). Tn*6944* was identified in an additional 156 *C. difficile* strains (78.0% – 100.0% NI), 97 of which belonged to clade C2 (**Table 5**). All *tet44*-positive *C. difficile* harboured Tn*6164* (80.3% – 100.0% NI), a 100 kbp genomic island containing *tet44* and *ant*(6)-Ib, a streptomycin resistance determinant (30).

**Figure 4.**
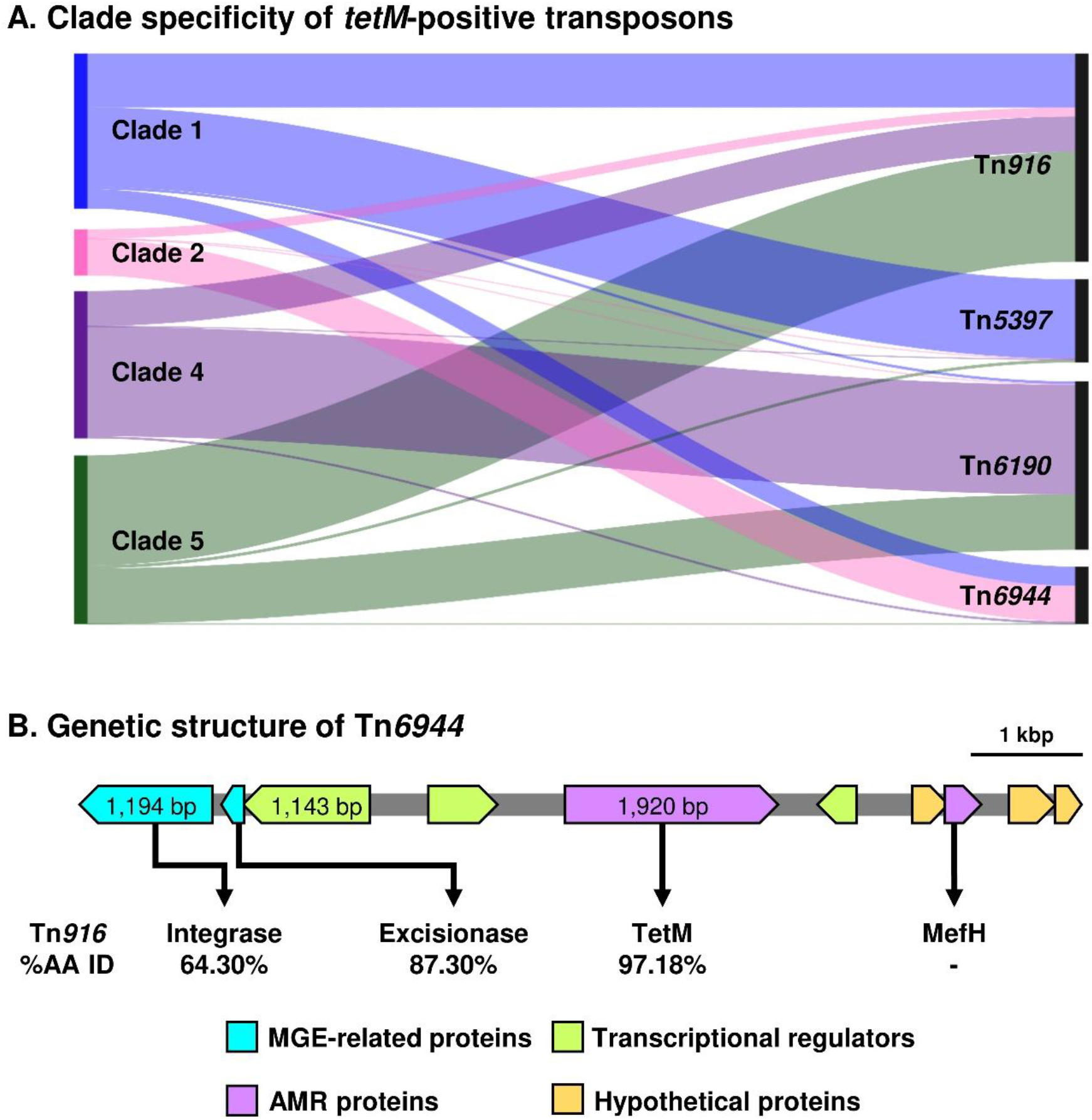
Clade specificity of *tetM*-positive transposons in *C. difficile*. **(A)** Sankey diagram shows the prevalence of four *tetM*-positive transposons commonly found in *C. difficile*. The left and right axes represent *C. difficile* clades and the transposons, respectively. The height of the left axis corresponds to the number of *tetM*-positive *C. difficile* strains in each clade, excluding strains with unknown transposons (clade 1, n = 419; clade 2, n = 208; clade 4, n = 711; clade 4, n = 688). **(B**) The genetic structure of the novel *tetM*-positive Tn, Tn*6944* [BK013348]. The amino acid sequences of the key elements in this transposon were compared to the elements found in Tn*916* [U09422.1].

### Vancomycin resistance

A complete *vanB* operon (*vanR*_*B*_, *vanS*_*B*_, *vanY*_*B*_, *vanW, vanH*_*B*_, *vanB* and *vanX*_*B*_ genes) was identified in one *C. difficile* strain, belonging to ST 11 (clade C5). This *vanB* operon was previously described to be phenotypically silent due to a ~2.1 kbp disruption of the *vanR*_*B*_ gene which is a response regulator and part of a key two-component system (31, 32). This strain was thus considered susceptible to vancomycin.

### Metronidazole resistance

SRST2 with the customised pCD-METRO plasmid database correctly identified the plasmid in 14 *C. difficile* genomes from the Boekhoud *et al*. study (25) (nine belonged to ST 15 and five belonged to ST 2). Apart from these strains, the pCD-METRO plasmid was found in only one *C. difficile* strain belonging to ST15 (clade C1, RT 010, non-toxigenic), the same RT reported in the Boekhoud *et al*. study (25). In total, only 10 of 223 *C. difficile* ST 15 strains (4.5%) contained the pCD-METRO plasmid.

### Rifamycin resistance

Points mutation in *rpoB* were found in 688 *C. difficile* strains (6.7%), with the highest prevalence in clade C4 (32.1%), followed by C2 (16.8%). The most common substitution was Arg505Lys found in 94.0% of the resistant strains, followed by His502Asn (49.4%), with 306 strains (44.5% of the resistant strains) having both substitutions. Besides rifamycins, a Gln1073Arg substitution in RpoB was also reported to be associated with reduced susceptibility to fidaxomicin (27). This substitution was not detected in this dataset.

### Carbapenem resistance

A total of 643 *C. difficile* strains (6.2%) had substitutions in either Pbp1 or Pbp3 conferring imipenem resistance, with the prevalence slightly higher in clades C2 and C4 (21.6% and 19.4%, respectively, p = 0.2786) than the other clades (collectively 1.4%, p<0.0001); 504 *C. difficile* strains had a substitution in Pbp1 (492 having A555T and 12 having L543H), 125 strains had a Y721S substitution in Pbp3 and 12 strains from ST 37 (C4) had substitutions on both Pbp1 (all A555T) and Pbp3.

In addition to the detection of point substitutions, carbapenemase-encoding genes were identified in two *C. difficile* strains; an unnamed strain [accession ERR2703875; ST 2, C1] carried SHV-1 and CD72 [accession SRR5367248; ST 81, C4] carried PER-1. By NCBI BLAST approach, the SHV-1 encoding gene was found on an element resembling a *Klebsiella pneumoniae* plasmid tig00001208_pilon [CP036443.1, 99.7% sequence identity, 35% coverage] and the PER-1 encoding gene was found on an element resembling *Acinetobacter haemolyticus* plasmid pAHTJR1 [CP038010.1, 99.8% sequence identity, 5% coverage].

### Other resistance types

Genotypic resistance determinants for five other antimicrobials were also identified. First, 124 *C. difficile* strains (1.2%) were positive for the *cfrB* gene which confers linezolid resistance (33). Resistance determinants for trimethoprim were identified in 147 (1.4%) *C. difficile* strains, six of which also harboured sulphonamide resistance determinants. Ninety-eight *C. difficile* strains (1.0%) carried chloramphenicol resistance determinants. The most common determinant was *catP* (92 strains, 93.9%).

In addition to the *C. difficile* class D β-lactamases which confer intrinsic cephalosporin resistance in *C. difficile* (17), a few *C. difficile* strains also had other classes of β-lactamases. Forty-three *C. difficile* strains carried genes encoding extended-spectrum β-lactamases (ESBL), the most common type belonging to the TEM family (36 strains), and five strains carried AmpC β-lactamase genes.

Finally, 1,250 *C. difficile* strains (12.1%) carried various aminoglycoside-resistance determinants. The most common determinants were *aac6-aph2* (666 strains, 6.5%), *aph-III* (279 strains, 2.7%) and *sat4* (271 strains, 2.6%) genes. Notably, 270 strains carried a locus containing *aph-III* and *sat4* genes adjacent to one another, 184 (68.2%) of which belonged to clade C5 (183 ST 11 strains and one ST 163 strain). This locus had 99.91% nucleic acid identity to a gene cluster found in *Erysipelothrix rhusiopathiae*, as described in a previous study (34).

## Discussion

The success of several epidemic *C. difficile* strains is thought to be associated with an AMR phenotype which provides a survival advantage for these *C. difficile* strains in the presence of antimicrobials while imposing little fitness cost (35–37). Resistance to several antimicrobial classes has been associated with specific *C. difficile* lineages: fluoroquinolone and rifamycin resistance and *C. difficile* ST 1 (C2) (10, 12), tetracycline resistance and *C. difficile* ST 11 (C5) (11), as well as resistance to various antimicrobial classes and MDR and *C. difficile* ST 37 (C4) (8). This study provides genotypic evidence to support these associations, demonstrated by the higher resistance prevalence and, especially in the case of tetracycline resistance in *C. difficile* ST 11, a higher diversity of resistance determinants in the associated clades.

Although the metadata was not complete (only 60.3% of strains had information on geographical origin and there was inadequate information on host species), some interesting findings can be seen in this genome subset. **Figure 3** demonstrates the difference in AMR prevalence in different continents which may reflect the use of antimicrobials in these regions. The most prominent example is fluoroquinolones which are strictly regulated in Australia and New Zealand but widely used elsewhere (38). Consequently, there was a stark difference in the prevalence of FQR between Australia and the other three regions. Besides fluoroquinolones, the high prevalence of MLS_B_ and tetracycline resistance, especially in Asia, is suggestive of the overuse of these antimicrobials in the region (39).

Based on a large sample size, which should give an accurate representation of the *C. difficile* population, this study provides a global atlas of genotypic AMR determinants in *C. difficile*. In general, one resistance determinant appeared to dominate in most antimicrobial classes. For example, *ermB* and *tetM* genes were found in almost 90% of *C. difficile* strains with genotypic resistance to MLS_B_ and tetracycline, respectively. Fluoroquinolone and rifamycin resistance was also mainly determined by a single substitution in GyrA (Thr82Ile) and RpoB (Arg505Lys), respectively. This is similar to other Gram-positive bacteria, such as *Staphylococcus aureus* (40), where one genotypic determinant is responsible for a resistance phenotype in a majority of the bacterial population and is in contrast to many Gram-negative bacteria, such as several members in the *Enterobacteriaceae* (41), where resistance to an antimicrobial class can be conferred by several genotypic determinants. The dominance of a single genotypic determinant accommodates the development of genotype-based rapid detection kits for drug-resistant *C. difficile*, similar to real-time PCR assays for methicillin-resistant *S. aureus* (42). Such tools can be beneficial for surveillance for *C. difficile* outbreaks in the future.

Another benefit of a large sample size and next-generation sequencing (NGS) is the power to detect rare genotypic determinants. The most notable finding was the detection of carbapenemase-encoding genes in two *C. difficile* strains, STs 2 and 81, comprising approximately 0.02% of the population. Previously, carbapenem resistance in *C. difficile* has been mainly associated with point substitutions on Pbp1 and Pbp3 which cannot be transferred horizontally and only confer imipenem resistance (26). On the contrary, many carbapenemases provide resistance to a wide range of carbapenem antimicrobials and are capable of horizontal transfer (43). The detection of carbapenemase-encoding genes is concerning, as *C. difficile* mainly resides in the colon, the same habitat as many pathogenic *Enterobacteriaceae*, and transfer of these genes could give rise to carbapenem-resistant *Enterobacteriaceae* (CRE), another urgent threat in AMR (3). Conversely, *C. difficile* can also serve as a reservoir of these resistance genes. Indeed, the gene encoding SHV-1, one of the carbapenemases found in this study, was found on an element similar to a *K. pneumoniae* plasmid (tig00001208, GenBank accession CP036443.1; 99.7% NI), suggesting a possible inter-phylum transfer event between these two organisms, although this plasmid was classified as an IncF plasmid according to PlasmidFinder (44). Generally, the host range for IncF plasmids is limited to only within the Family *Enterobacteriaceae* (45). Further study is thus needed to confirm that this horizontal transfer is possible.

Recently, two novel resistance determinants for MLS_B_ antimicrobials were found in Asian *C. difficile* isolates; *erm*(52) and *mefH* (21). In a larger population of *C. difficile*, these two genes were found in 1.4 – 1.5% of *C. difficile* strains, approximately six times more prevalent than *ermG*, a gene previously believed to be the second most prevalent resistance determinant in *C. difficile* (8). Failing to detect these two determinants could partially explain the discrepancy between resistance genotype and phenotype in earlier studies (23). Indeed, the inclusion of *erm*(52) improved the concordance between clindamycin resistance genotype and high-level clindamycin resistance phenotype to 100% and *mefH* provided concordant genotype to *C. difficile* strains with isolated erythromycin resistance (21). Further characterisation of *mefH* revealed that the gene was located adjacent to *tetM* on a newly defined transposon Tn*6944* (**Figure 4B**). This transposon has also escaped detection and characterisation despite being present mainly in ST 1 (clade C2), a strain that has been extensively studied (10, 46). Interestingly, even though tetracycline resistance was a key factor in the evolution of the epidemic *C. difficile* ST 11 due to its use in agricultural practices (11), this antimicrobial was not included in the antimicrobial susceptibility panel in a pan-European study (47, 48). Tetracycline resistance was also never mentioned in studies involving *C. difficile* ST 1, perhaps because the prevalence in this lineage was much lower than that of FQR mutations (7.1% vs 82.3%, respectively).

As an obligate anaerobe, *C. difficile* is intrinsically resistant to aminoglycosides. Additional resistance determinants to these antimicrobials are not beneficial to the bacterium and unlikely to be conserved in the genome. Thus, the presence of aminoglycoside resistance determinants should reflect recent, and likely continuous, inter-species gene transfer with taxa in diverse environments such as the animal gut and soils. The most common aminoglycoside resistance determinant was *aac6-aph2*, a bifunctional gene found in *Staphylococcus* spp. and *Enterococcus* spp. (49), commensal species commonly found in the human and animal gut. Interestingly, many ST 11 (C5) strains also carried an *aph-III* and *sat4* cluster, a gene cluster found in *E. rhusiopathiae* which inhabits the porcine gut (50), supporting the animal origin and One Health importance of this lineage (34). Indeed, aminoglycosides have been heavily used in both agricultural and veterinary practices (51). The presence of aminoglycoside resistance determinants in *C. difficile* highlights another aspect of AMR in *C. difficile*; the role of *C. difficile* as a reservoir of AMR genes. Aminoglycosides remain a key treatment option for serious staphylococcal and enterococcal infections, such as infective endocarditis, in conjunction with β-lactams antimicrobials (52). Resistance to aminoglycosides in these pathogens complicates treatment of these infections which may result in adverse clinical outcomes. Thus, colonisation with *C. difficile* carrying these resistance determinants may pose an additional risk of treatment failure in these patients.

This study utilised the direct analysis of raw sequence reads without the need for genome assembly which enabled the characterisation of a large dataset within a relatively short time (approximately 5 min of CPU time [16 cores] per strain as opposed to more than 30 min of CPU time per strain for a *de novo* assembly pipeline). SRST2 provides rapid MLST and AMR genotyping (15). SRST2-based AMR genotyping can be performed using three types of databases: well-characterised databases of accessory AMR genes (19, 20, 22), species-specific gene allele databases (e.g., the PubMLST database), as well as customised databases. The latter was used in a previous study on a smaller dataset, the results of which were similar to a standard approach using BLAST on annotated draft genomes (53).

Besides the lack of complete metadata, another limitation of this study was the lack of comparative phenotypic data, as the study was performed on a publicly-available genome dataset. However, many key AMR genotypes were reported to have a high correlation with phenotypic characteristics (23, 53). Thus, the prevalence values reported in this study should reflect the resistance prevalence in *C. difficile* population. Also, this study only reports the presence or absence of genotypic AMR determinants and does not take into account the different alleles of the genes, as the alleles were not included in the databases used in the analyses (19, 20). Further analyses on the allelic distribution across *C. difficile* population may provide additional information on the spread of AMR genes.

In conclusion, almost half of *C. difficile* strains studied carried at least one genotypic resistant determinant. The resistance prevalence was higher among clades C2, C4 and C5 which have been associated with epidemic *C. difficile* STs 1, 37 and 11, respectively. Though resistance to antimicrobials for treatment of CDI is rare, this study provides evidence to support the role of AMR in the spread of *C. difficile*, as well as the role of *C. difficile* as a reservoir of accessory AMR genes, most notably aminoglycoside resistance determinants and carbapenemase-encoding genes.

## Acknowledgements

This work was supported, in part, by funding from The Raine Medical Research Foundation (RPG002-19) and a Fellowship from the National Health and Medical Research Council (APP1138257) awarded to D.R.K. K.I. is a recipient of the Mahidol Scholarship from Mahidol University, Thailand. This research used the facilities and services of the Pawsey Supercomputing Centre [Perth, Western Australia].

## Additional information

The **Supplementary Data** is available at 10.6084/m9.figshare.14623533.

